# The genetic architecture of neurodevelopmental disorders

**DOI:** 10.1101/009449

**Authors:** Kevin J. Mitchell

**Affiliations:** Institute of Genetics and Institute of Neuroscience, Trinity College Dublin, Dublin 2, Ireland

**Author notes:** To appear in: The Genetics of Neurodevelopmental Disorders (Wiley) Edited by Kevin J. Mitchell http://eu.wiley.com/WileyCDA/WileyTitle/productCd-1118524888.html.

## Abstract

Neurodevelopmental disorders include rare conditions caused by identified single mutations, such as Fragile X, Down and Angelman syndromes, and much more common clinical categories such as autism, epilepsy and schizophrenia. These common conditions are all highly heritable but their genetics is considered to be “complex”. In fact, this sharp dichotomy in genetic architecture between rare and common disorders may be largely artificial. On the one hand, much of the apparent complexity in the genetics of common disorders may derive from underlying genetic heterogeneity, which has remained obscure until recently. On the other hand, even for supposedly Mendelian conditions, the relationship between single mutations and clinical phenotypes is rarely simple. The categories of monogenic and complex disorders may therefore merge across a continuum, with some mutations being strongly associated with specific syndromes and others having a more variable outcome, modified by the presence of additional genetic variants.

## Introduction

There are several hundred known genetic syndromes that affect neural development and result in intellectual disability (ID), epilepsy or other neurological or psychiatric symptoms. These include recognised syndromes that often manifest with symptoms of autism spectrum disorders (ASD) or schizophrenia (SZ), such as Fragile X syndrome, Rett syndrome, tuberous sclerosis, velocardio-facial syndrome and many others. For ASD, it has been known for many years that these syndromes account for a significant but still small fraction (5–10%) of all cases (Miles, 2011). What has not been clear is whether such cases, associated with single mutations, represent a typical mode by which such conditions arise or are, alternatively, exceptional and quite distinct from the general etiology of idiopathic ASD, epilepsy, SZ or ID (Wray and Visscher, 2010). Other common disorders including dyslexia, specific language impairment, obessive-compulsive disorder, etc., will not be considered here in detail, though the general principles probably apply.

In general, the genetic architecture of common NDDs has been considered to be “complex” or multifactorial (Plomin et al., 2009; Sullivan et al., 2003). This is usually taken to mean that many causal factors, both genetic and non-genetic, are involved in each affected individual. Under this view, the large group of currently idiopathic cases have a very different genetic architecture from the small number of known monogenic cases. An alternative view is that the vast majority of cases of these conditions are caused by independent mutations in any one of a very large number of genes. According to this model, these diagnostic categories of idiopathic cases represent artificial groupings reflecting our current ignorance, rather than natural kinds.

Here, I consider the theoretical underpinnings and empirical evidence relating to the genetic architecture of NDDs. These have been greatly influenced by technological advancements which have allowed various types of genetic variation to be assayed. Studies over the past several years have revealed an extreme level of genetic heterogenity and complexity for common NDDs, with the discovery of high-risk mutations in a large number of single loci and additional complexities in the causal architecture in individuals.

## Theoretical considerations

Linkage studies have clearly shown that common NDDs are not caused by mutations in one particular gene, leading to the unchallenged conclusion that variants at many loci must be involved *across the population* (e.g., (O’Rourke et al., 1982; Szatmari, 1999)). However, models of the genetic architecture of these conditions differ in two additional, independent parameters: (i) the number of variants thought to contribute to disease *in any individual* (from one or a few to many, possibly thousands), and (ii) the presumed frequency of risk alleles (from very rare to very common). The differences between models have profound implications for finding causal variants, predicting disease risk, discovering underlying biology and developing treatments for particular patients. More fundamentally, they represent very different ways of conceptualising these conditions.

## Number of causal alleles per individual

At one extreme, a model of Mendelian inheritance with genetic heterogeneity proposes that each case is caused by a single mutation, but that these can occur in any one of a large number of different loci (McClellan and King, 2010; Mitchell, 2011; Wright and Hastie, 2001). The types of mutations could include chromosomal aberrations that change the copy number of multiple genes, or mutations affecting a single gene. This model also encompasses diverse modes of inheritance, from *de novo* mutations to dominant or recessive inheritance. Fundamentally, this model conceives of common clinical categories such as SZ, ASD, epilepsy, ID, etc., as umbrella terms for large numbers of distinct conditions that happen to manifest with similar symptoms (Betancur, 2011; McClellan et al., 2007; Mitchell, 2012; Mitchell and Porteous, 2011; Ropers, 2008).

There are many precedents for this kind of genetic heterogeneity, including the genetics of congenital deafness (Lenz and Avraham, 2011), various forms of blindness, such as retinitis pigmentosa (Wright et al., 2010) and the many known Mendelian forms of intellectual disability (Ellison et al., 2013) and epilepsy (Poduri and Lowenstein, 2011). What differs with these conditions is that they typically display clear-cut Mendelian modes of inheritance, which is rarely the case for NDDs.

Moreover, linkage studies have been highly successful in identifying causal loci involved in specific Mendelian sub-types of these disorders, whereas they have produced highly inconsistent findings for common diagnostic categories, such as ASD and SZ (see below). Partly due to the failure of linkage studies to zero in on specific causal loci, an alternative model of polygenic inheritance became the dominant paradigm in the field (Risch, 1990).

The polygenic model proposes that common disorders arise from the combined action of a large number of risk alleles in each affected individual (Falconer, 1965; Plomin et al., 2009). Regrettably, the term polygenic has been used more loosely in recent literature to refer simply to the involvement of many loci across the population, where the number of contributing loci per individual remains unknown and could be as low as one (Purcell et al., 2014; Sullivan et al., 2012). I use polygenic here in the original sense to refer to conditions caused by the combined effects of multiple variants per individual.

Under the polygenic model, many risk variants are floating through the population and their independent segregation generates a continuous distribution of risk variant burden. Individuals at the extreme end of this distribution are thought of as passing a threshold and consequently developing disease (Falconer, 1965) (Figure 1). This model views common disorders effectively as unitary conditions, reflecting a common etiology – people with disease are simply at the tail end of a single distribution that extends continuously across the whole population. The distribution in this case is of the imagined latent variable, “liability”, which is presumed to exist and to be normally distributed, but which cannot be measured directly. It can be translated, statistically, into the highly discontinuous distribution of observed risk (in relatives of affected individuals, for example), by invoking an essentially arbitrary threshold, above which disease results. This liability-threshold model is statistically convenient but highly abstract (Mitchell, 2012).

**Figure 1.**
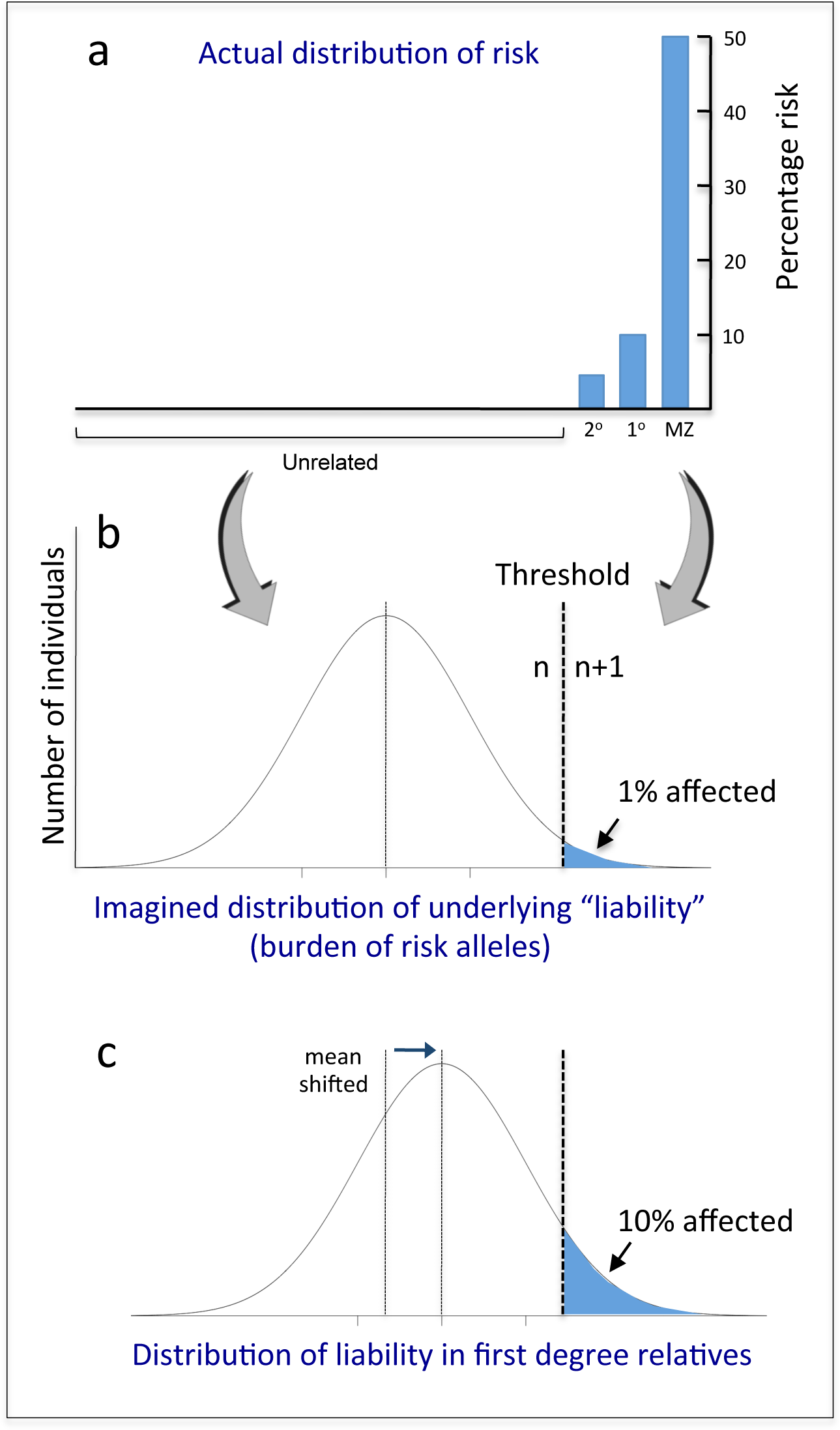
The liability-threshold model. A discontinuous distribution of observed risk across the population (a) is represented as reflecting an underlying latent variable, the “liability”, which is assumed to be normally distributed (b). A threshold of burden is invoked to regenerate the observed discontinuity. The mean liability of siblings of affected individuals is presumed to be shifted towards the threshold (c), explaining the greater disease incidence in this group compared to the population average. This yields a scenario analagous to response to selection for a quantitative trait, enabling heritability to be estimated (Falconer, 1965). Reproduced, with permission, from (Mitchell, 2012)).

An extension of this model considers the disorder as arising from the extremes of a number of actual quantitative traits, or endophenotypes (Gottesman and Gould, 2003; Meyer-Lindenberg and Weinberger, 2006). Common neuropsychiatric conditions affect multiple cognitive or social functions or faculties, such as working memory, executive function, sociability, etc. All of these traits also show a distribution across the unaffected population and all show moderate heritability. This led to the suggestion that individuals diagnosed with conditions like ASD or SZ may simply be at the extreme end of the normal distributions for several of these traits at the same time (Gottesman and Gould, 2003; Meyer-Lindenberg and Weinberger, 2006).

The corollary of that idea is that the genetic variants contributing to variation in such traits across the normal population *will be* the risk variants for such disorders. The hope was that such traits might have simpler genetic architectures than clinical diagnoses or at least that any genetic associations would be more obvious, as these traits reflect functions supposedly closer to the action of the genes.

## Frequency of risk alleles – evolutionary considerations

In addition to the number of loci involved, the frequency of each causal allele in the population is an independent parameter of models of genetic architecture. Polygenic models could involve rare or common alleles, or a mixture of both. The common disease/common variants (CD/CV) model proposes that common diseases arise from the cumulative burden of a number of common risk variants that float through the population, or at least that some of the causal variants would be common (Reich and Lander, 2001).

When applied to NDDs, a major problem arises for the CD/CV hypothesis. Such disorders significantly reduce fitness, with early onset, higher than average mortality and much lower than average fecundity (Keller and Miller, 2006). The CD/CV hypothesis must therefore address how genetic variants that predispose to the disorder could become common in the population in the face of negative selection (Keller and Miller, 2006). Various explanations have been invoked, including different forms of balancing selection, where the disease-causing variants are beneficial in another context. They could, for instance, increase fitness in a subset of individuals with a different genomic context, i.e., those who do not develop disease but carry some of the risk variants. Or it could be that such risk variants were beneficial in a different environment, such as our species’ recent past. There is, however, no evidence to support either of these contentions (Keller and Miller, 2006), and examples of balancing selection remain exceptional (Mayo, 2007; Olson, 2012).

An alternative explanation is that in situations where the effects on risk of each common variant are very small, and only expressed in a minority of carriers for any one variant, they are effectively invisible to selection. This may well apply under a model involving a huge number of loci with infinitesimal effect sizes. It could also arise if common alleles act as modifiers of rare mutations, but have no effect in most carriers. On the other hand, given a large effective population size, even a small average decrease in fitness across all carriers of a genetic variant means that natural selection can quite effectively keep its frequency low (Agarwala et al., 2013; Eyre-Walker, 2010; Gazave et al., 2013).

By contrast, a model involving multiple rare variants/mutations is completely congruent with evolutionary genetics as it explicitly incorporates an important role for natural selection in keeping the frequency of individual disease-causing variants low or even rapidly eliminating them. New variants constantly arise through *de novo* mutation, generating a balance between mutation and selection and maintaining the disorder at a certain prevalence in the population. The prevalence of a disorder then largely depends on the size of the mutational target – the number of genes that can be mutated which result in that particular phenotype (Keller and Miller, 2006; Rodriguez-Murillo et al., 2012).

The distinction between the two models is thus quite stark – on the one hand, the polygenic, CD/CV model implicates a standing pool of common, ancient variants floating through the population (Plomin et al., 2009). By contrast, the model of genetic heterogeneity involving rare mutations (McClellan and King, 2010) is consistent with a much more dynamic spectrum of human genetic variation, with causal mutations winking in and out of existence, some being immediately selected against, others persisting for several generations (Lupski et al., 2011; Olson, 2012). Under this model, the more recent and thus rarer variants would have a larger phenotypic effect, though necessarily in fewer individuals. More severe conditions should be characterised by a higher contribution from *de novo* or recent alleles, while those where the effects of fitness are lower could involve a greater contribution from less rare (possibly even common) alleles, which could persist in the population for longer (Agarwala et al., 2013; Eyre-Walker, 2010; Simons et al., 2014).

This model fits with recent data showing the extent of rare variation in human populations and the frequency distribution of deleterious alleles (Abecasis et al., 2012; Gravel et al., 2011; Keinan and Clark, 2012; MacArthur et al., 2012). Rare alleles collectively make up 90% of the variation across the population. There is, moreover, a strong skew towards rarer, more recent alleles among those predicted to deleteriously affect a protein (including nonsense mutations, frameshifts and those affecting splicing particularly) (Keinan and Clark, 2012; Kryukov et al., 2007). This implies that such alleles tend to be under strong negative selection and, conversely, that alleles with large biological effects tend to be rare. Because *de novo* mutations have not yet been subject to negative selection, they are likely to include the most highly penetrant alleles.

The descriptions above represent the extreme versions of these two models. As we will see below, the empirical evidence actually favours an integrative model for the genetic architecture of NDDs. This encompasses a heterogeneity of causal architectures across individual cases, with some being more genetically complex than others. It also combines effects of multiple variants in individuals to explain observed complexities in relating genotypes to phenotypes. This model applies not just to common clinical categories but also to rare, identified syndromes, where phenotypic variability and genetic modifier effects are becoming more apparent.

## Empirical evidence

### Familiality

Several characteristics of the observed familiality of common disorders have been taken as evidence against a model of simple Mendelian inheritance with genetic heterogeneity and in favour of a polygenic burden model of inheritance.

1. With rare exceptions, most families do not show an obviously Mendelian pattern of inheritance – these disorders are characterised by familial *aggregation*, rather than consistent patterns of *segregation*.
2. There is a high rate of sporadic cases – most affected children have normal parents and no affected first-degree relatives.
3. Recurrence risk increases with the number of affected children in a family.
4. Recurrence risk to siblings typically increases with severity of the defect in the proband.
5. Risk is greater when both parents are affected.
6. Risk to relatives falls off sharply with increasing degree of relationship to an affected proband.

All of these observations are consistent with the idea of an increased burden of risk alleles in some families, which would be indicated by both increased number of affected individuals and increased clinical severity and which would manifest as increased risk to subsequent children.

However, these observations are also consistent with a scenario where (i) many cases are caused by *de novo* mutations, explaining the high incidence of sporadic cases and rapid fall off in risk with increasing genetic distance, and, (ii) many causal mutations are incompletely penetrant for any particular clinical category. More highly penetrant mutations segregating in a family would lead to greater severity and a greater proportion of individuals reaching the criteria for a clinical diagnosis. The observed patterns of familiality thus do not distinguish between models of genetic heterogeneity and polygenic burden (Mitchell and Porteous, 2011).

In fact, the association of increased risk to siblings with increasing severity in the proband likely does not hold for all NDDs. The relative risk to siblings of patients with intellectual disability is paradoxically much lower (no higher than population average in fact) if their relative has severe intellectual disability than if they are only mildly affected (Roberts, 1952). This is consisent with a scenario where mutations causing intellectual disability with high penetrance are effectively immediately selected against and thus must arise *de novo*, while those causing only mild impairment are far more likely to be inherited.

### Linkage studies

Linkage studies for specific rare syndromes have been highly successful in identifying causal loci. Examples include Rett syndrome, tuberous sclerosis, Hirschsprung’s disease and many others (e.g., (Amir et al., 1999; Escayg et al., 2000; Luo et al., 1993; Wan et al., 1999)). In these cases, the fact that they were discrete conditions was recognised *a priori* on the basis of typical symptom clusters, thus permitting the grouping of patients from different families.

By contrast, linkage studies based on common, broader clinical diagnostic categories were not so successful. Given the scarcity of large pedigrees with multiple affecteds, it was necessary to pool samples from large numbers of smaller families in the hopes of identifying common loci. Though many linkage peaks were reported, these were often not replicated in subsequent studies and generally did not lead to the identification of specific genes.

These results, along with segregation analyses, clearly rule out mutations in one or a small number of specific loci as causing the majority of cases of any common NDD. The inconsistency of linkage results for common NDDs such as SZ has been given as evidence in favour of a polygenic model of inheritance (Risch, 1990). However, negative linkage results are also fully expected under a model of extreme genetic heterogeneity (Agarwala et al., 2013; Mitchell and Porteous, 2011) and thus do not distinguish between models.

### Endophenotypes

The endophenotype model for the genetic architecture of NDDs predicts that the mean phenotypic value of unaffected relatives of patients should be shifted towards the extreme end of the distribution of the endophenotype trait in question. This does seem to be the case for some endophenotypes, though not for all. For example, relatives of patients with SZ show mean values for some psychological measures that are lower than the population average, falling between the means of patients and controls (Allen et al., 2009; Braff et al., 2008). This trend extends to certain motor abilities and sensory processing measures and even various brain imaging measures.

What is not clear from those studies is whether this represents a consistent shift across all relatives or an effect seen in only a subset. The latter scenario seems to hold for ASD, where only a subset of relatives display what has been termed the Broad Autism Phenotype, scoring above a threshold on measures of autistic-like traits. For example, the BAP was apparent in 14–23% of parents of autistic children, compared to 5–9% of parents from a community sample (Sasson et al., 2013), with the remainder scoring in the normal range.

This more bimodal distribution of effects in relatives is consistent with a model of causation by rare mutations, with incomplete penetrance. Many relatives would not carry the causal mutation and would thus not differ from controls. Others would carry the mutation without developing the full clinical condition, but could show more subtle effects. This has been observed in clinically “unaffected” carriers of many pathogenic CNVs, for example (Stefansson et al., 2014). Alternatively, in cases caused by two or more mutations, relatives might carry only one of those and thus show a lesser effect (Berg and Geschwind, 2012; Girirajan et al., 2012) (Figure 2). The fact that the values of some endophenotypes are altered in some relatives thus does not distinguish between models of genetic architecture.

**Figure 2.**
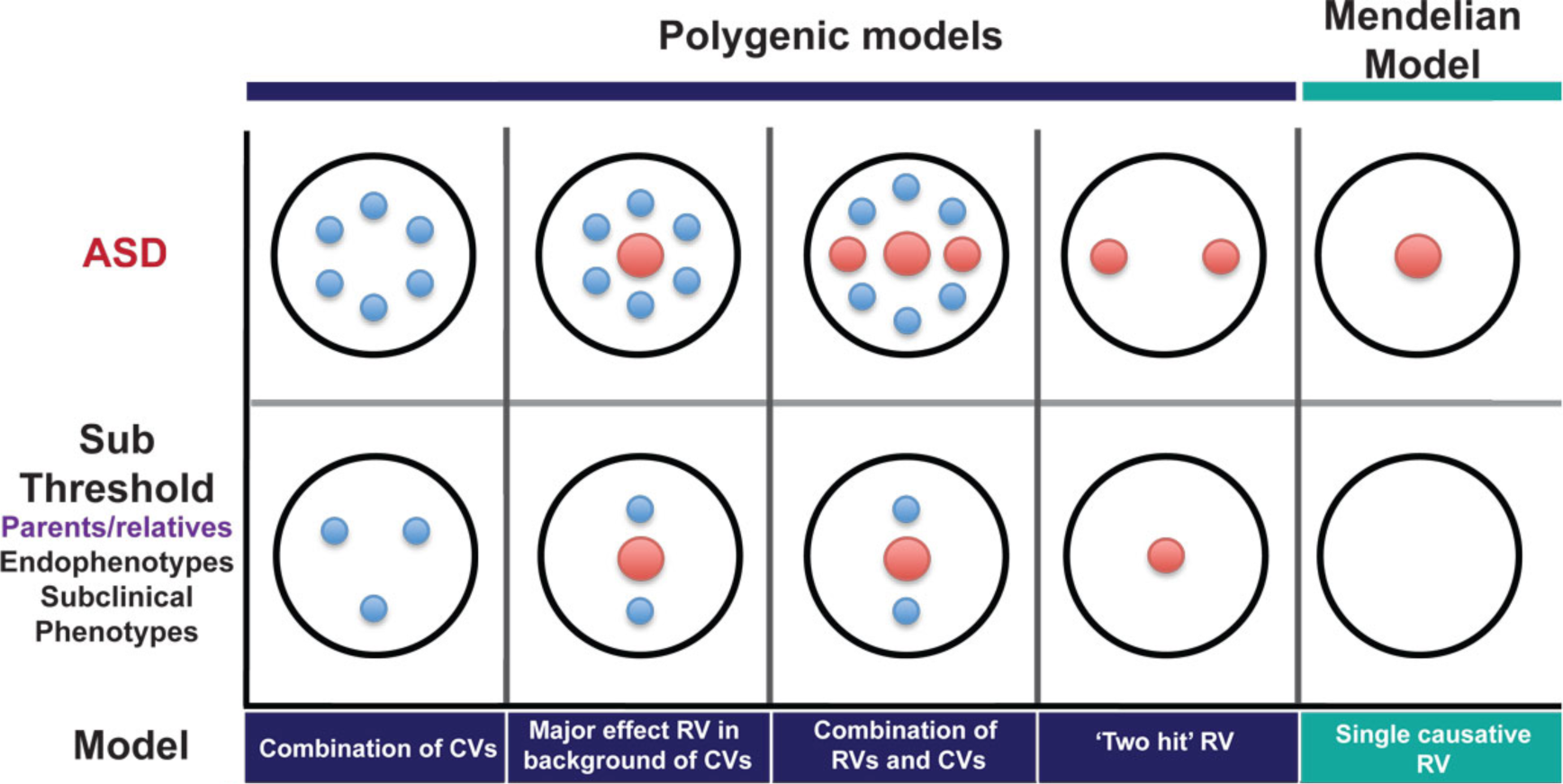
Expectations of risk allele burden and endophenotypes in relatives under a range of models of genetic architecture. Large red circles represent high-risk mutations, small blue circles represent common variants. The top row shows possible causal architectures for patients with ASD. The bottom row shows the expected distributions of causal variants in clinically unaffected relatives for each of these scenarios. (Reproduced, with permission, from (Berg and Geschwind, 2012)).

In a related vein, studies of the heritability of autistic-like traits across the general population have been taken by some as arguing that the genetics of these traits generally overlaps with the genetics of ASD. These studies have found the heritability is about the same at the extremes of the normal distribution of these traits, where patients with ASD diagnoses tend to score, as in the middle (Lundstrom et al., 2012; Robinson et al., 2011).

By itself, this does not prove, or even really argue for, a model whereby patients with ASD are those who fall at the extreme end of a unitary population distribution. The phenotypic values of ASD patients on those traits may fall at that position of the distribution for a different reason. If we consider an analogy with height, for example, it is clear that the genetics of dwarfism or giantism are quite distinct from the genetics of the normal distribution. A similar situation holds for the genetics of severe intellectual disability, which is clearly distinct from the genetics of IQ generally.

In addition, many single mutations are highly pleiotropic, affecting multiple endophenotypes at once, even though the genetics of such traits across the general population are largely non-overlapping. Overall, there is thus little support for the model that clinical patients with diverse symptoms happen to lie at the extreme end of the normal distributions of multiple independent traits.

### Common variants – genome-wide association studies

Direct tests of the hypothesis that common variants contribute to risk of disease were made possible by the development of the human Haplotype Map (Consortium, 2003), which enabled genome-wide association studies (GWAS) (Hardy and Singleton, 2009). These studies assay the frequencies of different alleles at hundreds of thousands of single-nucleotide polymorphisms (SNPs), distributed across the genome. These are positions where two alternative DNA bases are both at high frequency in the population. They reflect an ancient mutation that has spread to some extent throughout the population, typically due to genetic drift. There are tens of millions of such sites across the genome, but, due to uneven patterns of recombination across the genome, many such SNPs fall into haplotype blocks that tend to be co-inherited. As a result, sampling hundreds of thousands of SNPs, defined by the HapMap Project (Consortium, 2003), allows one to assay common variants across a much larger proportion of the genome.

The idea behind GWAS is very simple: if a common variant increases risk of disease, then the frequency of that variant should be higher in cases with the disease than in healthy controls (Hardy and Singleton, 2009; Risch and Merikangas, 1996). So, if a SNP shows that pattern, then either that SNP, or a variant that tends to be co-inherited with it, can be said to be associated with greater statistical risk of the disease. The problem is, if that statistical increase in risk is very small, then it requires a massive sample to detect it. This problem is greatly exacerbated by the statistical burden of correcting for all the multiple tests performed when assaying hundreds of thousands of SNPs at once.

Initial GWAS for NDDs, such as SZ, epilepsy and ASD, revealed no genome-wide significant hits (Anney et al., 2010; Kasperaviciute et al., 2010; Need et al., 2009). The sample sizes of these studies were relatively small but large enough to exclude the existence of any common variants with even a modest statistical effect on risk (increased risk of 2-fold or more). Somewhat larger studies for SZ have identified statistical associations with a number of common SNPs, with quite small effect sizes (odds ratios of <1.2) (Purcell et al., 2009; Shi et al., 2009; Stefansson et al., 2009). Along with additional loci implicated in larger studies, these collectively account for ∼3% of the total genetic variance affecting disease risk (Purcell et al., 2009; Ripke et al., 2013). At the time of writing, results from even larger GWAS for SZ have been reported, though not yet published. These mention over 100 associated SNPs, with even smaller individual effect sizes, though the overall genetic variance explained has not increased from earlier studies (Wright, 2014).

Recognising the etiological overlap between diagnostic categories, a recent study conducted a cross-disorder GWAS, encompassing cases with ASD, SZ, ADHD, bipolar disorder and major depression. Four loci gave genome-wide significant hits and seven others approached this level. Some signals were associated with single disorders, but most gave signal across disorders (Consortium, 2013). Again, the effect sizes were very small and the overall variance explained was less than 3%.

GWAS have also been conducted for a number of clinical or psychlogical endophenotypes. A small number of statistically signficant hits have been found (Alliey-Rodriguez et al., 2011; Connolly et al., 2013; Goodbourn et al., 2014; Knowles et al., 2014). These should be interpreted with caution, however, as they derive from small samples, have not been replicated and test for association with multiple traits at once. GWAS with larger samples, looking at individual dimensions of clinical symptoms, have not detected any hits at genome-wide significance (Bramon et al., 2014; Fanous et al., 2012). In addition, a large number of candidate gene associations with diverse endophenotypes have been reported in the literature. These have typically not held up well in subsequent replication attempts and the vast majority likely represent false positives (Flint and Munafo, 2013; Ioannidis et al., 2011).

Given the lack of variance in disease liability explained by currently identified SNPs and the possibility that studies to date have simply been underpowered, it is interesting to ask more generally, how much variance could theoretically be explained by common alleles collectively? A new quantitative genetics technique, which does not rely on individual SNPs reaching genome-wide significance, has been applied to GWAS results to attempt to estimate this quantity (Yang et al., 2010; Yang et al., 2011). This method of genome-wide complex trait analysis (GCTA) looks for a signature of increased (but still distant) relatedness amongst cases, compared to that amongst controls, and uses such a signature to estimate heritability. Estimates from this method for the overall percentage of genetic variance that is tagged by common SNPs are quite high for SZ and ASD (23% and 40–60%, respectively) (Klei et al., 2012; Lee et al., 2012). However, confidence in these figures is undermined by questions about the methodology and underlying assumptions of this technique and the interpretation of the results (Browning and Browning, 2011; Lee et al., 2012). In particular, the idea that a hypothetical, minuscule increase in risk could be detected in people cryptically related at only the 4th or 5th cousin level, when the increase in risk for 1st cousins is only about 2-fold for SZ and ASD (Lichtenstein et al., 2006; Sandin et al., 2014), seems to warrant some skepticism. Moreover, despite claims that this method indicates a large collective role for many common variants, it actually cannot distinguish either the number of loci involved or the frequency of causal alleles (discussed in more detail in Box 1).

#### Box 1. Estimating the overall contribution of common variants

Recently, a new type of analysis of GWAS data has been developed that purports to estimate how large a contribution common alleles could collectively make to quantitative traits, including the modeled liability to complex disorders (Lee et al., 2011; Yang et al., 2011). Genome-wide complex trait analysis, or GCTA, analyses SNP data from case-control datasets, but, rather than looking for signatures of association with individual SNPs, it uses these data merely to estimate genetic similarity between ostensibly unrelated pairs of individuals. This can then be correlated with phenotypic similarity, where continuous traits are concerned, to estimate how heritable the trait is. The logic for continuous traits is thus the same as with twin or family studies – the comparison is just carried out over much larger genetic distances, with correspondingly smaller phenotypic similarity.

For dichotomous traits, such as disease diagnosis, however, the logic is inverted from that of twin studies. Here, you start with people with a certain degree of phenotypic similarity (they all have the disease) and ask if they have higher genetic similarity (to each other than to a set of controls). A signature of increased mean genetic similarity across all pairs of cases is taken as evidence of heritability of risk for the disorder over large genetic distances. According to this method, a quantitative value for the percent of genetic variance tagged by common SNPs can be derived from the similarity matrix. The application of this method to SZ datasets has led to the assertion that 23% of the variance in liability to SZ is captured by SNPs and that a substantial proportion of this variation must be the result of common causal variants (Lee et al., 2012). A similar analysis for cases diagnosed with ASD concludes that “common genetic polymorphisms exert substantial additive genetic effects on ASD liability” and estimate the magnitude of these effects as explaining between 40–60% of additive genetic variance (Klei et al., 2012).

These values are obviously very substantial, but how much confidence can we have in their estimation and interpretation? The values are extrapolated from a tiny signal of increased (but still very distant) genetic similarity among cases, compared to controls, raising a general concern that such a signal may reflect artefacts or noise. There are a number of methodological concerns with this approach, to do, for example, with the statistical corrections required to exclude effects of cryptic population stratification (Browning and Browning, 2011) and to correct for highly skewed ascertainment of cases and controls relative to the true population prevalence of the disorder (Lee et al., 2011).

More generally, recurrence risks for SZ and ASD decrease sharply with increasing genetic distance and are only on the order of 1.5 to 2-fold for first cousins (Lichtenstein et al., 2006; McGue et al., 1983; Sandin et al., 2014). It seems likely, therefore, that any increased risk to fourth or fifth cousins would be negligible. The idea that a statistical signature of such an effect, if it exists at all, could be detected, measured accurately and extrapolated to give a definitive value of variance tagged by common SNPs thus seems inherently questionable.

Even taking the data at face value, however, it is not possible to infer that these signals are driven by causal effects of common variants, as stated by the authors of one of these studies: “From the analyses we have performed, we cannot estimate a distribution of the allele frequency of causal variants” (Lee et al., 2012). Allele-sharing between distant relatives is often concentrated in one or two genomic segments derived from a common ancestor (Ralph and Coop, 2013), meaning that increased sharing of rare variants in such segments could explain the supposed tiny average increased risk of disease among distant relatives (or, conversely, increased distant relatedness among cases). Using common SNPs to estimate heritability across distant relatives thus simply does not inform on the number of causal variants in the population or in individuals or the frequency of causal alleles.

Returning to those SNPs that do show genome-wide significant hits, what do these statistical associations mean? First, they do not imply that the SNP that is assayed is necessarily the causal variant itself. Each SNP tags an extended haplotype with many other common variants, so that the GWAS signal only implicates a general locus in the genome as containing some variant (or variants) affecting risk. Moreover, from that signal alone, it is impossible to infer how common the causal variant is. Modeling suggests that some GWAS signals may tag rare variants at a locus, which may by chance be more associated with one haplotype over another (Chang and Keinan, 2012; Dickson et al., 2010; Wang et al., 2010). Others have countered that rare variants cannot explain GWAS signals (Wray et al., 2011), but simulations incorporating the important parameter of negative selection suggest that GWAS signals across a locus can indeed be quite consistent with the presence of multiple, rare causal variants at that locus in the population (Thornton et al., 2013).

Empirical studies, involving resequencing of GWAS loci, have now found several instances where GWAS signals for various disorders or traits can be partially or largely attributed to effects of rare variants at the associated locus (Oosterveer et al., 2013; Sanna et al., 2011; Saunders et al., 2014; Thun et al., 2013). This effect has not been seen in all cases, however (Hunt et al., 2013). Furthermore, the general level of consistency of direction of allelic associations across distant populations, though by no means universal (Ntzani et al., 2012), is somewhat higher than expected under a model of synthetic associations as the sole drivers of GWAS signals (Carlson et al., 2013; Marigorta and Navarro, 2013). Overall, it thus seems likely that at least some of the reported GWAS signals for many disorders reflect a functional role for associated common variants.

This leads to the question of how to interpret the effect sizes of associated SNPs in GWAS. These are usually expressed as odds ratios, which summarise the statistical increase in relative frequency of one SNP allele in cases versus controls. For disorders that are not very common, this approximates the relative risk – the increased likelihood of being a case, given the presence of the risk allele. Most odds ratios from GWAS are in the range of 1.05 to 1.2-fold increased risk. How can this statistical effect across the population be related to biological effects in individuals?

The most straightforward possibility is that everyone who carries that risk allele is at very slightly higher risk of developing disease than those who carry the alternate allele. An alternative interpretation is that that average signal reflects a much more potent effect, but in far fewer people. Such a situation could arise where: (i) rare variants of larger effect fall predominantly on that common haplotype, i.e., the signal is driven by synthetic associations, as described above, or, (ii) a common allele acts as a strong genetic modifier of particular rare mutations, at the same or different loci, but has essentially no effect in most individuals, who do not carry such mutations. Examples of such modifiers will be discussed below.

### Rare mutations – copy number variants

Many rare neurodevelopmental syndromes (such as Down, Williams, Angelman, Prader-Willi syndromes and many others) are associated with specific chromosomal anomalies, including deletions or duplications of sections of chromosomes (also known as copy number variants, as they change the number of copies of genes within the deleted or duplicated segment). These conditions were initially distinguished by the consistent clustering of behavioural and non-psychological symptoms, such as typical facial morphology, for example. The causal chromosomal anomalies were discovered by classical cytogenetics and subsequently defined molecularly.

The development of array technologies for detecting CNVs across the genome allowed these efforts to become far more systematic and powerful (Sebat et al., 2004). The application of these technologies and the realisation that CNVs could also be detected using SNP arrays led to the discovery of numerous additional CNVs that are associated with increased incidence of various common NDDs (e.g., (Cooper et al., 2011; Kirov et al., 2009; Marshall et al., 2008; Mefford et al., 2010; Sebat et al., 2007; Walsh et al., 2008); reviewed in (Cook and Scherer, 2008; Grayton et al., 2012; Merikangas et al., 2009)). The risk associated with such CNVs is recognisable because they recur at a low but detectable frequency at particular sites in the genome, due to local properties of genomic organisation (Liu et al., 2012). It is thus possible to find many people with effectively the same chromosomal deletion or duplication and assess rates of illness in this group.

Though individually rare, the CNVs so far identified can collectively account for a significant proportion of previously idiopathic cases of conditions like ASD (>10%) and SZ (>5%). One of the most striking findings from this work has been the lack of respect for clinical diagnostic boundaries in the effects of such CNVs. The same CNVs have been detected in patients with ASD, SZ, epilepsy, ADHD, ID and other clinical presentations (Cook and Scherer, 2008; Grayton et al., 2012; Merikangas et al., 2009). The genetic etiology of these conditions is thus clearly overlapping (Coe et al., 2012; Craddock and Owen, 2010; Moreno-De-Luca et al., 2013), a finding that is reinforced by large-scale epidemiological studies and by analyses of mutations in single genes (see Chapter 2).

### Single-gene mutations

CNVs delete or duplicate chunks of chromosomes and typically affect more than one gene. But NDDs can also be caused by mutations in single genes, which are now also being discovered at an increasing rate, thanks to the development of next generation sequencing technologies. In addition to those associated with syndromic forms of mental illness, such as Fragile X syndrome and Rett syndrome, early studies had identified a small number of single-gene mutations associated mainly with psychiatric manifestations in particular families. These include DISC1, where carriers in a large Scottish pedigree of a translocation that disrupts the gene manifest with a variety of psychiatric diagnoses (Millar et al., 2000), and genes encoding neuroligin-3 and neuroligin-4, mutations in which were found in families with multiple individuals affected by ASD (Jamain et al., 2003).

As with the initially identified chromosomal syndromes, some argued that these might be isolated examples that are not relevant to the majority of idiopathic cases. This idea has turned out to be untenable, as more and more cases associated with single-gene mutations are discovered. Next-generation sequencing studies, using both family and case-control designs, have identified numerous point mutations, or single-nucleotide variants (SNVs), associated with high risk of NDDs (Allen et al., 2013; Chahrour et al., 2012; Cukier et al., 2014; Fromer et al., 2014; Iossifov et al., 2012; Lim et al., 2013; Neale et al., 2012; O’Roak et al., 2011; O’Roak et al., 2012; Piton et al., 2010; Purcell et al., 2014; Sanders et al., 2012; Xu et al., 2011; Yu et al., 2013). As with CNVs, most of these mutations are associated with diverse clinical manifestations. Mutations in any one gene are individually very rare, as expected, but an overall excess of damaging SNVs in patients with NDDs compared to controls suggests that a large portion of the burden of disease may be accounted for by such rare mutations collectively (Allen et al., 2013; Cukier et al., 2014; Fromer et al., 2014; Kenny et al., 2013; Purcell et al., 2014).

One of the most important findings from studies of CNVs and SNVs is that a significant proportion of the pathogenic mutations arise *de novo*, in the generation of sperm or eggs, rather than being inherited from a carrier parent (Ku et al., 2012) (Chapter 3). Typically, mutations that have higher penetrance for more severe phenotypes will be more likely to have arisen *de novo* than to have been inherited, as carriers are less likely to have children. Current estimates suggest that as many as 50% of cases of ASD may be attributable to *de novo* mutations (Ronemus et al., 2014). This is likely to be even higher for severe forms of ID (Vissers et al., 2010), but lower for later-onset disorders with smaller effects on fitness, such as SZ and bipolar disorder.

This finding has several general implications. First, it illustrates the general point that even common NDDs can be caused by single, dominant mutations. Second, it shows that such conditions can be genetic but not inherited, reconciling high hertiability (based on MZ twin concordance) with the high level of sporadic cases. Finally, it further undermines the quantitative genetics framework, which is premised on the idea of a standing pool of variation that simply gets shuffled around from generation to generation.

As more and more high-risk mutations are identified, more and more cases will move from the idiopathic pool to the pool with known high-risk mutations (Figure 3). However, while some such mutations will define new syndromes, it would be a mistake to think of causality generally in such simple terms. The incomplete penetrance and variable phenotypic expressivity of many single mutations, whether *de novo* or inherited, suggests additional layers of complexity in relating genotypes to phenotypes.

**Figure 3.**
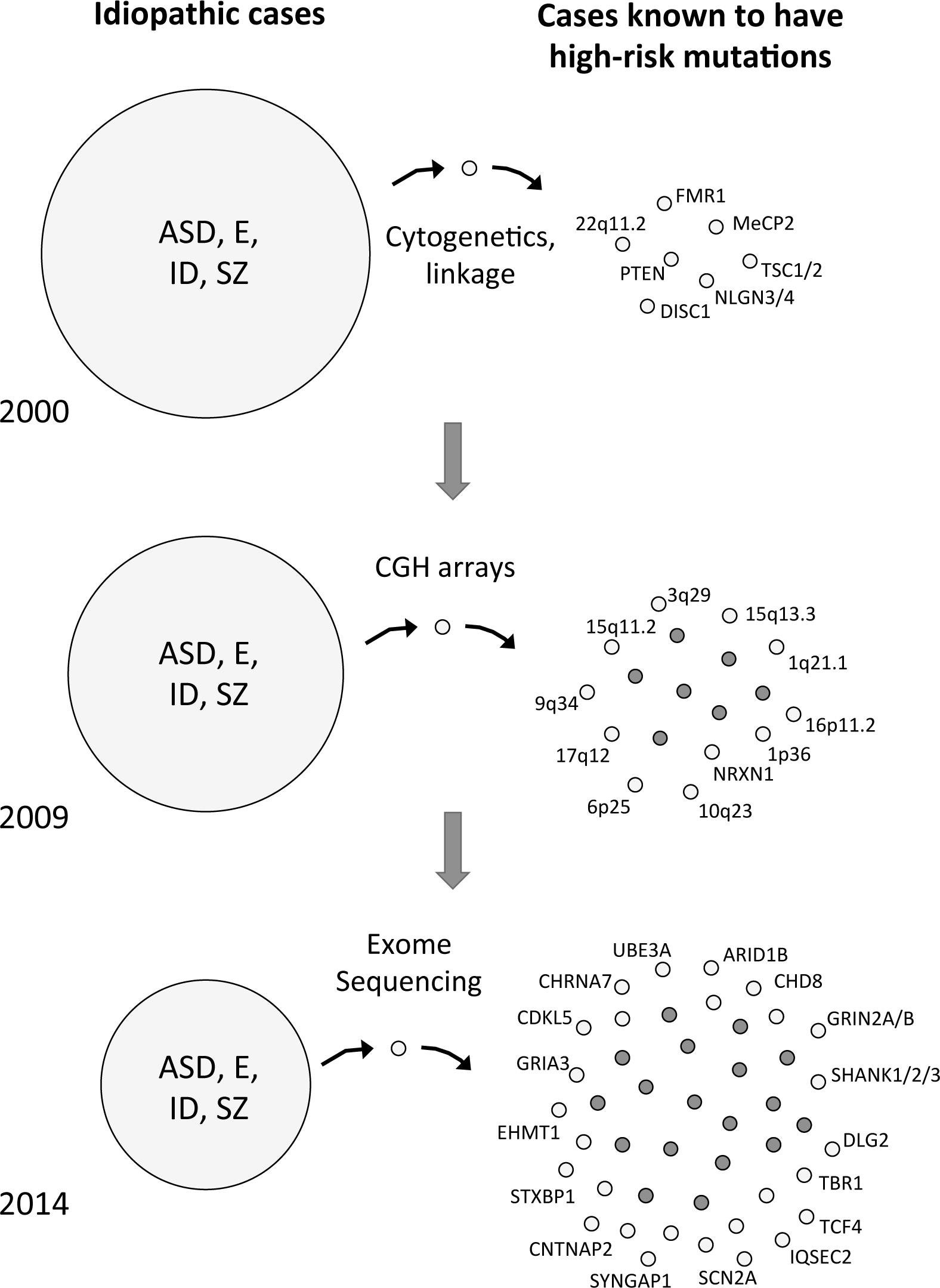
The cumulative identification of genetic causes of neurodevelopmental disorders. The circle on the left represents the current pool of idiopathic cases, reflecting the level of ignorance at the time. The small circles on the right represent cases carrying rare, high-risk mutations. New technologies including comparative genome hybridisation (CGH) arrays and next-generation sequencing of exomes or genomes have allowed a continuing stream of discoveries of new risk mutations (lighter circles), thus shrinking the pool of idiopathic cases. Note that only an arbitrary set of examples of such mutations are shown; the real list runs to many hundreds.

## Complex genotype-phenotype relationships

### Incomplete penetrance and variable expressivity

While the list of high-risk mutations is growing all the time, it is also clear that simple models relating genotypes at single loci to clinical phenotypes generally do not hold. The phenotypic expression of most such mutations is quite variable and penetrance for any specific diagnosis is typically incomplete. Of course, penetrance can be defined in other ways, which do not rely on reified clinical categories. For example, while the penetrance for many CNVs for SZ is relatively low (Vassos et al., 2010), the penetrance for a broader category including ASD and developmental delay is much higher (Kirov et al., 2014). In addition, while such CNVs are also detected at reduced frequency in healthy controls, a recent study has found that many are associated with general decreases in cognitive ability, even in clinically “unaffected” carriers (Stefansson et al., 2014).

Variability in phenotypic expression is also now becoming apparent even for mutations associated with specific syndromes, such as VCFS, Williams syndrome, Angelman syndrome and others. Prospective screening of psychiatric patients without syndromic diagnoses has revealed that the CNVs causing these syndromes also are found in patients with idiopathic symptoms of autism, epilepsy or other neurological or psychiatric manifestations (Grayton et al., 2012). The initial, narrow definition of specific syndromes thus likely reflects an ascertainment bias, in that discovery of these mutations was based on grouping together those patients with the most recognisably similar pattern of symptoms. A similar situation is observed for mutations causing inborn errors of metabolism. Though these are typically recognised due to their phenotypic effects in young infants, many such mutations are now also being implicated in adult-onset psychiatric patients with no previous diagnosis (Kayser, 2008; Sedel, 2012).

Another important factor in relating mutations in specific genes to specific clinical outcomes is allelic heterogeneity. Not all mutations at a particular gene will alter protein production or function in the same way. This is classically exemplified by mutations in different parts of the dystrophin gene, which cause the clinically distinct conditions of Duchenne or Becker muscular dystrophy. Many genes show a similar diversity of outcomes associated with mutations in different regions of the gene (Walsh and Engle, 2010). In addition, some mutations associated with severe cortical malformations or other developmental syndromes when homozygous, have now been found in heteroyzgous condition in less severely affected patients, manifesting mainly with psychiatric symptoms (Walsh and Engle, 2010).

### Genetic modifiers and oligogenic effects

The effects of primary mutations are commonly modified by genetic background. This is a truism in experimental genetics with model organisms, where strain background effects are commonplace – almost ubiquitous, in fact (Mackay, 2009; Nadeau, 2001; Phillips, 2008; Spiezio et al., 2012). The phenotypic effects of many mutations vary – sometimes hugely – between strains of mice or flies, for example. This has several interesting implications: first, and most obviously, the phenotype in individuals is often determined by more than one genetic variant. Second, some of the variants involved have little or no phenotypic effect alone (in cases where the phenotype in question does not vary between two strains in the absence of a major mutation, for example). Third, the existence of such cryptic genetic variation is evidence that the developmental system is capable of buffering substantial genetic variation without altering the phenotype (Gibson and Dworkin, 2004; Wagner, 2007). The latent effects of such variation may be released, however, in the presence of a serious mutation.

This is also true in human genetics. Many mutations associated with distinct Mendelian conditions are strongly modified by additional genetic variants (Badano and Katsanis, 2002; Cooper et al., 2013; Dipple and McCabe, 2000). This is true even for conditions associated with mutations in a single gene, such as sickle cell anemia, cystic fibrosis and Huntington’s disease, where severity, age of onset and progression can all be modified by specific variants in other genes. The manifestation of various NDDs also depends on background variants, as in Rett syndrome (Grillo et al., 2013; Renieri et al., 2003), Dravet syndrome (Singh et al., 2009) and Kallmann syndrome (Shaw et al., 2011), for example. Specific modifying variants have been identified for many genetic conditions (Cooper et al., 2013). Some of the modifying variants are themselves rare, but common variants can often make important contributions, significantly modifying the risk of specific mutations.

This scenario is exemplified by Hirschsprung’s disease, a neurodevelopmental disorder affecting the enteric nervous system (Alves et al., 2013). Rare mutations in 18 genes have been associated with this condition, including the RET and NRG1 genes. Importantly, common variants in both those genes also increase risk and are much more frequent in affected carriers of the rare mutations than in unaffected carriers. However, in the absence of a rare mutation, these common variants have little or no phenotypic consequence. These effects thus exemplify epistatic, or non-additive genetic interactions in determining individual phenotypes (Chapter 4).

To date, no specific modifying mutations have been definitively identified for more common NDDs, but this may reflect a lag in discovery, exacerbated by the higher level of primary genetic heterogeneity. It seems quite possible that some of the common variants identified by GWAS may be acting in this fashion – that is, the small statistical effects associated with some common SNPs, when averaged across the population, could be due to much larger effects in only a subset of individuals carrying rare mutations. Interestingly, GWAS signals for common NDDs have shown up in a number of loci in which rare mutations are associated with specific syndromes with neurological and psychiatric symptoms, such as Pitt-Hopkins syndrome (TCF4) (Forrest et al., 2014), Timothy syndrome (CACNA1C) (Bhat et al., 2012) and cerebellar ataxia (SYNE1) (Consortium, 2013; Noreau et al., 2013). These GWAS signals could be due to synthetic associations, but could alternatively reflect a situation like that in observed in Hirschsprung’s disease, where common variants have strong modifying effects. It will likely be necessary to first define carriers of specific primary mutations before these kinds of specific modifying effects can be recognised.

One common variant that has been demonstrated to have a large effect on the phenotypic outcome associated with neurodevelopmental mutations is the Y chromosome. This is most evident in ASD, where males are much more commonly affected than females (a 4:1 ratio) (Ronemus et al., 2014), but can also be observed in sex differences in the rates of many NDDs, including ADHD, dyslexia, SZ and others (Cahill, 2006). Analyses of the spectrum of mutations in autistic patients reveals that affected females tend to have more severe mutations than affected males. This is true for both CNVs (which affect many more genes on average in females) (Levy et al., 2011) and SNVs (which include more potentially deleterious mutations in females) (Jacquemont et al., 2014). Importantly, these effects extend to broader categories of NDDs and are also seen in the previous generation. When the mutation is inherited, it is significantly more likely to come from the mother than the father (Jacquemont et al., 2014; Ronemus et al., 2014). This suggests that men who carried such a mutation were more severely affected and thus less likely to become fathers in the first place.

These findings indicate that it takes a more severe mutation to push a female brain into an autistic state, or, conversely, that males are more susceptible to the effects of such mutations. This sex difference could be due to the Y chromosome itself, through its known influences on brain development and connectivity (Gilmore et al., 2007; Ingalhalikar et al., 2014; McCarthy et al., 2012; Wu and Shah, 2011). Alternatively, it may be not the presence of the Y, but the lack of one X chromosome that is important – this may make male development intrinsically less robust to the effects of mutations anywhere in the genome. However, the facts that not all developmental disorders show this male bias, and that the bias is uneven for different NDDs, seem more consistent with a Y chromosome effect.

In addition to modifying effects of common variants, a growing number of cases of NDDs are now turning up with more than one severe rare mutation. This has been observed for CNVs, where affected individuals who have inherited a CNV with relatively low penetrance more often have a second, potentially pathogenic CNV elsewhere in the genome (Bassuk et al., 2013; Girirajan et al., 2012; Girirajan et al., 2010). Similar events have been observed for known pathogenic single-gene mutations with incomplete penetrance alone (Chilian et al., 2013; Leblond et al., 2012; Schaaf et al., 2011). In these cases, both CNVs or mutations are likely having an effect alone and these may combine, additively or non-additively, to cause frank disease. By contrast, mutations with higher penetrance tend to arise *de novo* and are not associated with an excess of secondary events (Girirajan et al., 2012).

### Non-genetic sources of variance

The fact that concordance for NDD diagnoses is not complete between MZ twins indicates the presence of additional sources of variance beyond overall genotype. These may include environmental risk factors but can also reflect an often neglected non-genetic source of variance, which is intrinsic developmental variation.

*Environmental factors*: Epidemiological studies have associated a number of environmental risk factors, such as maternal infection during gestation, pre-term delivery, obstetric complications and others with statistically significant increased risk for NDDs ((Hamlyn et al., 2013), Chapter 6). The odds ratios for each of these broad categories of risk factors are typically low (less than two-fold). However, such risks may be unevenly distributed across the population. In particular, pathogenic mutations could make the developing brain more susceptible to the effects of such environmental insults, leading to a greater effect in genetically vulnerable individuals. While plausible, and arguably more likely than a uniform effect, this kind of gene-by-environment interaction remains to be directly demonstrated for NDDs (Chapter 6).

*Intrinsic developmental variation*. The outcome of development is inherently variable, as evidenced by physical differences between isogenic organisms, including monozygotic twins, or even between the two sides of nominally symmetrical organisms (Leamy and Klingenberg, 2005). Such differences are also observed at the neuroanatomical level, as in agenesis of the corpus callosum, for example, where this structure may be absent in one twin and present in the other (Mitchell, 2007; Ruge and Newland, 1996; Wahlsten, 1989). On a finer scale, the effects of many mutations are played out at a cellular level in a probabilistic fashion across the developing brain, so that the pattern of abnormalities may vary widely from one brain to the next (as with mutations causing cortical heterotopia, to take an obvious example). Thus, even in non-pathogenic circumstances, by the time they emerge from the womb, the brains of monozygotic twins are already quite unique (Clarke, 2012; Mitchell, 2007).

Such effects could presumably contribute to differences in emergent psychological traits and neuropsychiatric disorders between MZ twins. For example, while the heritability of epilepsy is quite high, the heritability of the specific anatomical focus is much lower (Corey et al., 2011), likely reflecting such stochastic events in neurodevelopment. Intrinsic developmental variability may thus make a large contribution to the non-genetic variance observed for NDDs, which can be sizeable.

Behavioural genetics studies typically divide the sources of phenotypic variance into genetic variance, shared family environment effects and a third term, called the “non-shared environment” (Plomin and Daniels, 2011). This term mathematically accounts for the incomplete heritability of a trait or lack of full concordance between monozygotic twins, reflecting an additional non-genetic source of variance in the population generally (Turkheimer and Waldron, 2000). The phrase non-shared environment is somewhat regrettable, as it implies an origin outside the organism. This has often been interpreted as reflecting an important role for personal experiences, such as interactions with peers or teachers, which may help to differentiate MZ twins from each other (Harris, 1998; Plomin and Daniels, 2011). Given the strong and consistent evidence that family environment has little or no effect on phenotypic outcome for NDDs, there seems little reason to think that peer interactions or other, non-traumatic, personal experiences would make such an important contribution. Nor is there any reason to expect differential exposure to environmental toxins or other risk factors would be greater between MZ twins than between different families.

In fact, the non-shared environment term also encompasses: (i) measurement error or mis-classification (an important source of variance for behavioural traits or psychiatric diagnoses especially), and, (ii) chance, or, in this case, intrinsic developmental variation. The developmental program is quite robust and, under normal circumstances, strongly canalises development towards a species-typical outcome (Wagner, 2007). This idea is captured in the epigenetic landscape, a metaphor developed by Conrad Waddington, to conceptualise the probabilistic, but also canalised nature of development (Waddington, 1957). (This original usage of epigenetic derives from the Greek term “epigenesis”, meaning the emergence of the organism, and does not relate to the molecular genetic usage of the term, referring to mitotically stable chromatin states).

The developmental program is robust to small changes (as evidenced by the presence of cryptic genetic variation (Gibson and Dworkin, 2004)) but this robustness is challenged by severe mutations. These tend to not only decrease the probability of a species-typical phenotype, but also increase the phenotypic variance, making the outcome more susceptible to noise and stochastic events (Wagner, 2007; Yeo et al., 2007). Interestingly, the ability of a developing organism to buffer the effects of specific mutations may itself be a genetic trait, reflecting what we might call “genomic reserve”. The possiblity that such a trait reflects the general mutational load in the genetic background is explored in Chapter 5.

### Heterogeneity and complexity of causal factors in individuals

The preceding sections paint a picture of the etiology of NDDs that involves complexities on many levels. First, there is tremendous underlying locus and allelic heterogeneity. This reinforces the view that many broad clinical categories of common NDDs do not represent natural kinds, at least in terms of etiology. The distinction between cases whose symptoms are associated with known causes and the much larger group of idiopathic cases is simply an expression of current ignorance, not a reifying principle that justifies treating diagnoses of exclusion as natural kinds. This heterogeneity largely undermines the quantitative genetics framework (Mitchell, 2012), which lumps together patients with common diagnoses under the assumption of a shared and unitary etiology (Robinson et al., 2014).

Second, there is heterogeneity in modes of causality across individuals (Figure 4). In some cases, a primary mutation will be readily recognisable. In others, multiple mutations or modifying variants may be at play. Conditions that are more severe are more likely to have a greater proportion of cases caused by individual, recent mutations (often *de novo*), while those with less severe manifestations are more likely to have one or more inherited mutations and modifying variants.

**Figure 4.**
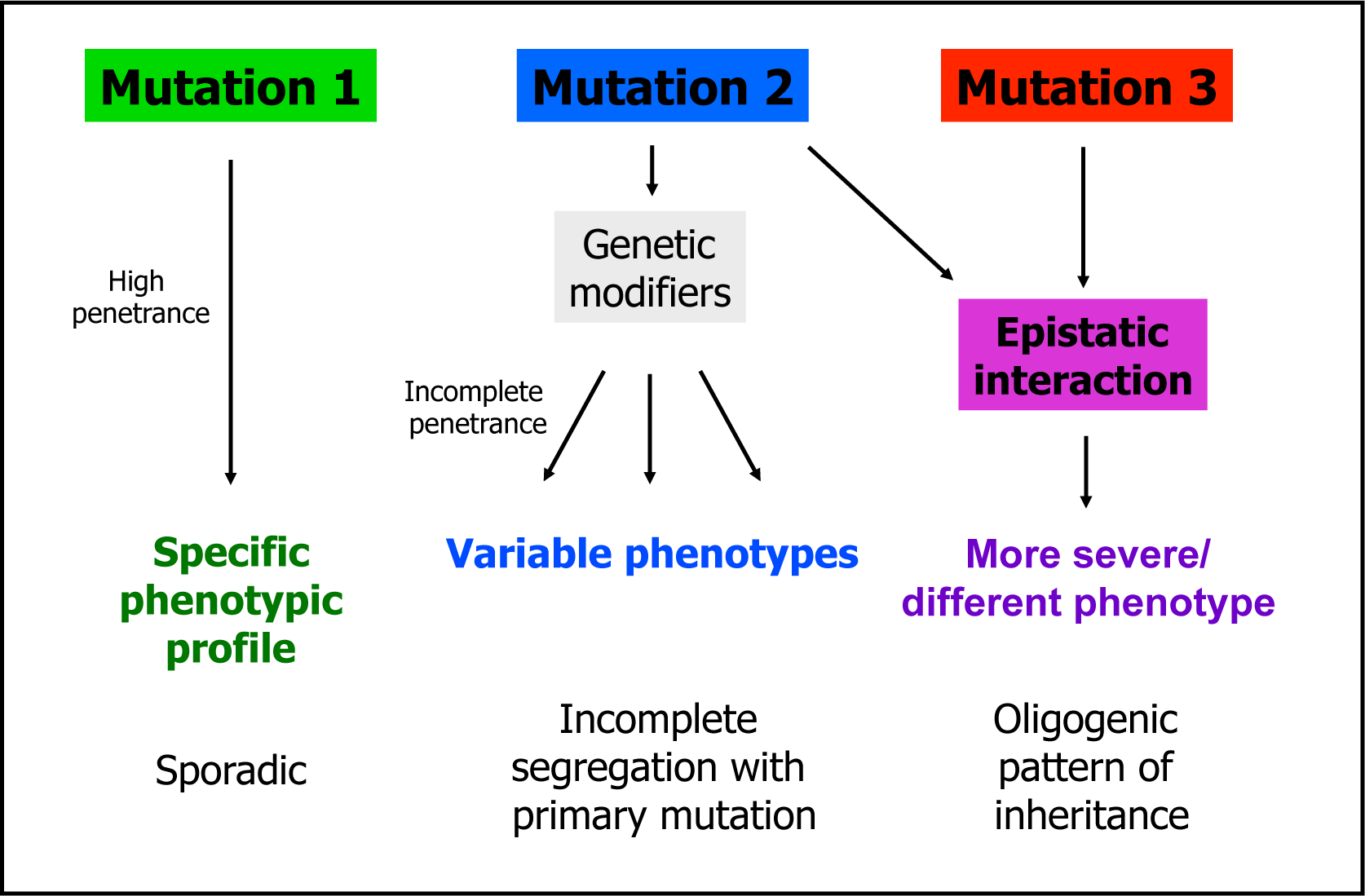
Heterogeneity of causal architectures across individuals. High penetrance mutations will most often be immediately selected against and so will typically arise *de novo*, rather than being inherited. Lower penetrance mutations will be more likely to be inherited and will often be modified by additional common or rare variants in the genetic background. Reproduced, with permission, from (Mitchell, 2011)).

Third, the sharp dichotomy between rare and common disorders has become blurred. On the one hand, we can now recognise within the broad categories of common disorders a growing number of distinct, very rare conditions (Figure 3). On the other, even the formerly recognised rare conditions show a previously unappreciated level of genetic complexity, with important contributions from genetic modifiers. Digenic or oligogenic causality, with a few contributing variants, is thus likely in many cases.

Whether the range of modes of causality extends to a highly polygenic architecture, involving a very large number of variants in some individuals, remains an open question. The statistical association of some variants of small effect across the population does not mean that they must act in this collective fashion. Rather than supposing a division whereby some cases are caused by mutations of large effect and others by the combined action of a very large number of mutations of small effect, an integrative model incorporates variants of low statistical effect size across the population (common or rare) as epistatic modifiers of major mutations (Mitchell and Porteous, 2011), as demonstrated for disorders like Hirschsprung’s disease.

The extreme level of locus heterogeneity is reflected in the diversity of functions of the genes implicated, which encode proteins acting in many different cellular processes. While many of the implicated genes encode proteins involved in neural development or synaptic function (discussed in detail in Chapters 7–9), others include chromatin regulatory proteins, basal translation factors, metabolic enzymes and miscellaneous proteins whose functions are not obviously neurodevelopmental. This raises an interesting and important question: how could disruption of so many different genes, with such diverse functions, lead to such similar outcomes – the states we recognise as ASD or SZ or epilepsy? An answer may lie in considering the symptoms of these disorders as emergent phenotypes.

### The genetics of emergent phenotypes

We already know of hundreds of different genes in which mutations can cause intellectual disability. This is perhaps not surprising, given the complexity of the human brain – it seems reasonable to expect that mutations in many different genes could impair its function. Somewhat more surprising is the idea that a similarly diverse set of mutations could give rise to the apparently much more specific phenotypes associated with common NDDs, such as autistic behaviour, psychosis, depression, hyperactivity or seizures. The symptoms of these conditions seem to reflect not just a decrement in function but the emergence of qualitatively novel brain states.

Understanding how mutations in so many diverse genes can give rise to these states involves the recognition that the relationship between the normal functions of genes and their resultant mutant phenotypes can be extremely indirect. This is especially true for phenotypes that reflect very high-level, emergent functions of complex systems. The kinds of cognitive processes impaired in ASD and SZ represent the highest-level functions of the human mind. These processes rely on the functions of a myriad distinct systems, each composed of multiple brain regions and fibre pathways, hundreds of cell types and thousands of gene products. Like “performance” of a fighter jet, high-level cognitive operations rely on the complex and dynamic interactions between all these components.

The fact that these systems are susceptible to mutations in many different genes is thus not so shocking. The upgrades to our cognitive hardware which arose through evolution may carry with them a certain vulnerability – the price of increasing complexity may be more ways to break down. We may in addition, as a social species, be highly attuned to notice subtle differences in function of the brain, which might be far less evident for other organs.

However, the fact that our neural systems tend to fail in particular ways, generating qualititatively novel brain states, remains an interesting puzzle. It seems likely, though, that this reflects organisational properties of the brain, rather than the functions of the perturbed genes. In particular, maladaptive reactivity of the brain to early differences may channel development towards discrete pathological states (Ben-Ari, 2008; Hulme et al., 2013; Lisman et al., 2008; Lodge and Grace, 2011).

The functions of the disease-associated genes are thus too diverse and too far removed from the emergent effects of the pathogenic mutations to think of them as “genes for” ASD, SZ or epilepsy. Nor is it accurate to conceive of them as genes for working memory or executive function or other high-level cognitive operations. The proximal effects of mutations in various genes, which arise at the molecular and cellular levels, will have cascading consequences over neural and cognitive development, with the phenotype of the organism sometimes being channeled by developmental systems and neural architectures to produce emergent states that we recognise as psychiatric or neurological conditions.

## Implications for research and clinical practice

The genetic architecture of NDDs is characterised by heterogeneity of causes across individuals and complexity of causes within individuals. This has a number of important implications for both research and for clinical practice:

1. Finding additional high-risk mutations by case-control comparisons will likely require very large samples, in order to distinguish the cuplrits from the innocent bystanders (Zuk et al., 2014). It may be possible and necessary to bootstrap our way from mutations that were strongly implicated under more specific study designs and to use biological knowledge to generate priors for inference of pathogenicity.
2. The identification of high-risk mutations in enough people may enable the secondary discovery of genetic modifiers. With increasing knowledge of such interactions, this may allow more accurate prediction of individual risks based on genome-types, not just single-mutation-genotypes.
3. Inferring genetic causality will likely remain a matter of probabilities. Nevertheless, as genetic information becomes available for more and more patients, it should be possible to discern which pieces of information are relevant for treatment (Box 2) (Chapter 13).
4. The segregation of patients based on genetic knowledge should greatly enhance the ability to define clinical subsyndromes (Bruining et al., 2014) and also to investigate the neurobiological phenotypes associated with specific mutations (Consortium, 2012; Stessman et al., 2014). It should be much more informative to characterise patients with the same mutation than to analyse patients grouped solely by broad clinical diagnosis, with high underlying heterogeneity.
5. The identification of high-risk mutations offers a proven discovery route to the underlying biological processes. Cellular and animal models with direct etiological validity, combined with our growing general understanding of how the brain works, should reveal pathogenic mechanisms and cascading pathways through which various mutations presumably converge on a narrower set of pathophysiological states (Arguello and Gogos, 2012; Mitchell et al., 2011) (Chapters 10, 11).
6. Combined, all these approaches offer the hope of rationally designing new therapies and intervention strategies based on a detailed understanding of the pathogenic and pathophysiological mechanisms in individual patients (Chapters 12, 14).

### Box 2. Causality and genetic diagnoses

Given the incomplete penetrance and variable expressivity of the known mutations implicated in NDDs, how should we think about genetic causality? In considering this issue, a clear distinction should be drawn between explanation versus prediction of illness, as the probability relationships are entirely different in these two directions.

In terms of predicting illness based on the presence of a known disease-associated mutation, the only information we have to go on is the penetrance of the mutation for the disorder in question. For the well known 22q11.2 deletion, for example, the risk of psychosis is about 30% (though the risk of any clinical diagnosis is much higher). By contrast, the rate of SZ in carriers of the NRXN1 deletion is about 6%. These increased risks may be deemed actionable in terms of reproductive decisions but would provide less justification, for example, for drastic preemptive clinical intervention in currently unaffected carriers.

On the other hand, the presence of these mutations in individuals who are *already affected* allows stronger inferences to be drawn about their contribution to illness. Here, one can compare the odds of someone having the illness, given the presence of the mutation, with the odds of them having the illness for some other reason (i.e., the population prevalence). For 22q11 deletions, the odds in favour of that deletion being a primary contributor to schizophrenic symptoms in that individual are thus around 30:1. For NRXN1 deletions, where prediction is quite weak, the inference of causality, given the illness having occurred, is considerably stronger: about 6:1 odds of the patient having the disease due to the NRXN1 mutation, as opposed to some other reason. This approach defines causality in a counterfactual rather than a reductive sense – it does not imply that the mutation in question was a sufficient cause, but does estimate the likelihood that it was a necessary cause in that patient.

However, patients with low penetrance mutations are more likely to also have a secondary mutation or additional genetic variants contributing to pathogenesis. This has been observed empirically (Girirajan et al., 2012) and fits generally with the known high concordance levels of MZ twins for common NDDs such as ASD and SZ. Simply put, most individuals who develop these diseases were at high risk of having done so. Though ascertainment biases likely inflate these figures somewhat (only seeing pairs condordant for illness, not for health), this does imply that most patients with these conditions carry high-risk *genome-types*. If the primary mutation is not potent enough on its own, this suggests the presence of additional accomplices.

The definition of discrete genetic syndromes and the assignment of categorical genetic diagnoses may be somewhat justified for high penetrance mutations but is thus less appropriate for patients with lower penetrance mutations. A more pragmatic approach will be simply to consider the potential relevance of any piece of genetic information to clinical management based on empirical observation, such as whether patients with mutations in Gene X tend to show particular symptoms or respond better to particular treatments.

Genetic information can thus be incorporated into clinical management without falling into the conceptual trap of issuing overly categorical genetic diagnoses (Chapter 13). In the future, the identification of modifying mutations, as for Hirschsprung’s disease for example, should make it possible to make more accurate predictions of risk based on an individual’s entire genome-type, and not just with a single mutation.

## Acknowledgments

Many thanks to David Goldstein and David Porteous for very thoughtful reviews of this chapter and to Dan Bradley and Svetlana Molchanova for helpful comments and conversations.

